# Performance evaluation of mesoscopic photoacoustic imaging

**DOI:** 10.1101/2022.10.17.512521

**Authors:** Lina Hacker, Emma L. Brown, Thierry L. Lefebvre, Paul W. Sweeney, Sarah E Bohndiek

## Abstract

Photoacoustic mesoscopy visualises vascular architecture and associated tissue structures at high resolution at up to 3 mm depth. The technique has shown promise in both preclinical and clinical imaging studies, with possible applications in oncology and dermatology, however, the accuracy and precision of photoacoustic mesoscopy has not been well established. Here, we present a performance evaluation of a commercial photoacoustic mesoscopy system for imaging vascular structures. Typical artefact types are first highlighted and limitations due to non-isotropic illumination and detection are evaluated with respect to rotation, angularity, and depth of the target. Then, using tailored phantoms and mouse models we demonstrate high system precision, with acceptable coefficients of variation (COV) between repeated scans (short term (1h): COV=1.2%; long term (25 days): COV=9.6%), from target repositioning (without: COV=1.2%, with: COV=4.1%), or from varying *in vivo* user experience (experienced: COV=15.9%, unexperienced: COV=20.2%). While our findings support the robustness of the technique, they also underscore the general challenges of limited field-of-view photoacoustic systems in accurately imaging vessel-like structures, thereby guiding users to correctly interpret biologically-relevant information.

## Introduction

Extending the range of macroscopic and microscopic photoacoustic imaging (PAI) approaches, raster-scanning photoacoustic mesoscopy has been recently introduced^1^ for visualization of tissue structures up to 2-3 mm depth^2,3^. The innovative nature of mesoscopy lies in combining wide-bandwidth, high-frequency acoustic detectors with fast nanosecond-pulsed laser excitation, enabling three-dimensional imaging of optically absorbing molecules such as haemoglobin and melanin^4,5^ at the mesoscopic scale (in-plane spatial resolution: ~20 μm)^2,3^. In the preclinical setting, photoacoustic mesoscopy has already been demonstrated for: longitudinal *in vivo* studies in several mouse models of cancer^6,7^; whole body imaging of zebrafish using a 360° multi-orientation approach^8^; and gastrointestinal imaging^9^. In clinical studies, it has shown particular potential in skin imaging, revealing individual skin layers, benign nevi^10,11^, hyperthermia effects^12^, or pathophysiological biomarkers of inflammatory skin diseases^2,13^.

Volumetric visualization of vasculature and skin layers is one of the key strengths and main applications of photoacoustic mesoscopy, however, the accuracy and precision for delineating targets of interest, such as vessel structures, has yet to be comprehensively evaluated. A well-known limitation of practical PAI systems lies in the limited apertures of their illumination and detection arrays, enabling only visualization of targets that are quasi-perpendicular to the direction of the transducer array^14^. Moreover, the non-isotropic illumination from the sides of the ultrasound detector array results in inefficient light delivery to the region of interest^15^, thereby leading to limited penetration depth, particularly in specimens that exhibit high absorption. Finally, the limited view can cause in-plane or out-of-plane artefacts, reducing the clarity of the images^16–19^. In addition to these technical limitations impacting accuracy, precision-related variations in image data arise from temporal, positioning, or operator-dependent factors. Detailed precision studies have been performed for other commercially available PAI systems, including a preclinical tomography system^20^ and a handheld clinical system^21,22^ which have established relevant bounds on the reliability and biological relevance of the extracted photoacoustic data. Such studies are crucial to ensure data reproducibility and accurate data interpretation as the application of mesoscopic PAI in preclinical and clinical research expands.

Here, we conduct a detailed technical validation of a commercial photoacoustic mesoscopy instrument to assess both accuracy and precision of the system and use our findings to identify common artefacts and suggest approaches to maximize image quality. Imaging limitations of photoacoustic mesoscopy are evaluated by characterising common artefact types and systematically analysing signal variations resulting from the restricted aperture of the illumination and detection array. Precision is then assessed both using tailored phantoms and *in vivo* using mouse models to account for system variation arising over time, target repositioning and user experience. We conclude by outlining recommendations for data acquisition. By presenting this validation study for photoacoustic mesoscopy systems, we hope to guide users with similar PAI setups, assisting them in system validation, data handling and data interpretation.

## Methods

### Mesoscopic photoacoustic image acquisition

The photoacoustic mesoscopy system (RSOM Explorer P50, iThera Medical GmbH, München, Germany; Figure 1A,B) has been described in detail elsewhere^29^. Briefly, laser light is generated by a 532-nm laser (pulses: 1 ns; ≤1 mJ/pulse) and delivered through a customized 2-arm fibre bundle (spot size: 3.5×5mm). Photoacoustic signals are detected by a spherically focused LiNbO_3_ detector (centre frequency: 50 MHz; bandwidth: 10-90 MHz; focal diameter: 3 mm; focal distance: 3 mm; f number: 1). The recorded data is amplified by a low noise amplifier of 63 dB gain. The scanning head is attached to two motorized stages and coupled to the sample surface by an interchangeable water-filled (2 mL) interface. For coupling of the object to the lower side of the interface, commercial ultrasound-gel (Aquasonic Clear, Parker Laboratories, Fairfield, NJ, USA) was used. The ultrasound gel was centrifuged to remove air bubbles and warmed before application. The interface was positioned on the object by moving the stage in x, y and z directions. Images were acquired over a field of view of ≤12 × 12 mm (step size, 20 μm). The acquisition of one image took approximately 7 min.

**Figure 1:**
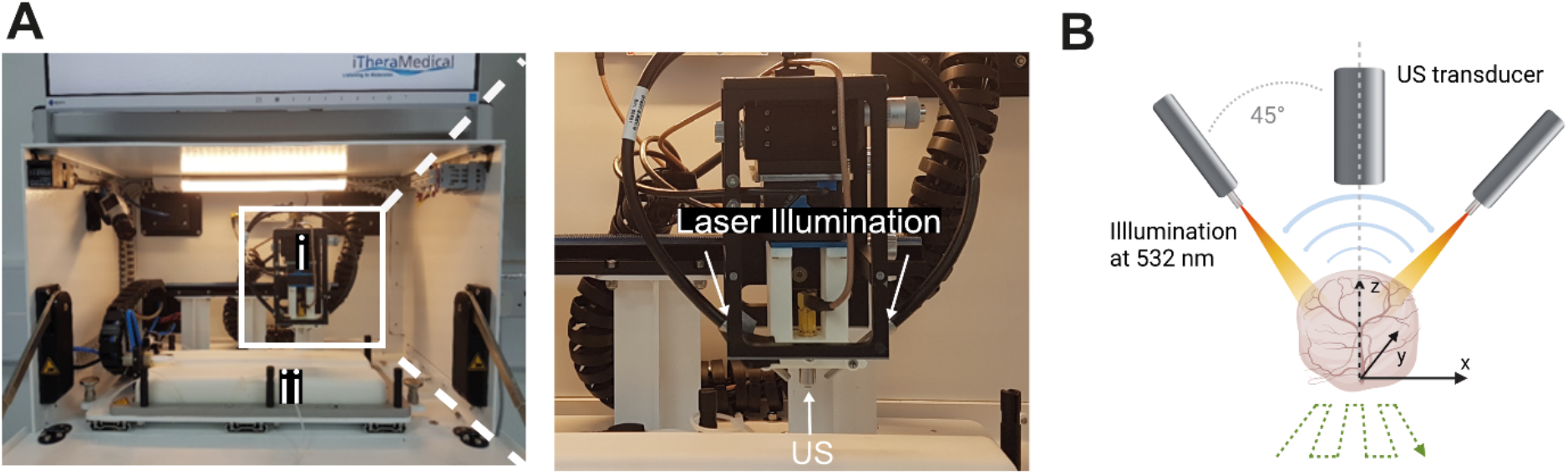
Overview of the photoacoustic mesoscopy system. A) Photograph of the system. (i) denotes the scan head and (ii) the heated mouse bed. The scan head is enlarged in the second photograph. Top arrows are pointing on the two illumination fibres, whilst the bottom arrow points on the ultrasound (US) transducer. B) Schematic illustrating the functioning of the system. Created with Biorender.

For phantom imaging, the phantom was placed underneath the transducer and aligned to the transducer direction in the position of interest. About 10 ml of ultrasound gel was used to couple the transducer surface to the phantom medium. Care was taken for the transducer interface not to touch the phantom surface.

For *in vivo* imaging, temporal variability with and without replacement was assessed over a time frame of 35 mins (n=5 images) with an n=5 per sample group. ‘With replacement’ is defined as full removal of the mouse from the heat pad including cleaning off the coupling ultrasound gel. For evaluation of the impact of operator experience, an operator with > 1 year experience in preclinical photoacoustic mesoscopy imaging was regarded as experienced, and an operator with < 10 days experience in preclinical photoacoustic mesoscopy imaging was regarded as inexperienced.

### Phantom preparation

#### Base material

For systematic evaluation of artefacts using geometric phantoms, agar was chosen as the bulk phantom material to provide structural support for imaging targets due to its facile and fast preparation method. A 1.5% tissue-mimicking^23^ agar mixture was prepared according to the protocol by Joseph et al^20^. Briefly, intralipid (2.08 v/v%; Merck, 68890-65-3) was used to mimic tissue-like scattering conditions and Nigrosin (0.62 v/v% of Nigrosin stock solution [0.5 mg/mL Nigrosin in deionised water, Merck, 8005-03-6]) was added to mimic tissue-like absorption.

For evaluation of long-term system precision, copolymer-in-oil base material was chosen^25^. The material was prepared according to Hacker et al^24^. Briefly, the phantom was composed of 30 w/v% high molecular weight SEBS (Sigma Aldrich 200557-250G) and 8 w/v% LDPE (Alfa Aesar 43949.30) in mineral oil (Sigma Aldrich-330779-1L), with 0.03 w/v% TiO_2_ (Sigma Aldrich 232033-100g) added to provide optical scattering and 0.0007 w/v% Nigrosin (Sigma Aldrich 211680-100G) added to provide optical absorption. The acoustic properties were characterised using a through transmission substitution system (available at NPL, London, UK)^24^, yielding a speed of sound of 1483.5 ± 0.17 m·s^−1^ and an acoustic attenuation of 6.73 ± 0.04 dB·cm^−1^ at 5 MHz. The optical properties were determined using a custom-built double-integrating sphere (DIS) system^24^, yielding a reduced scattering coefficient of 0.4 mm^−1^ and an optical absorption coefficient of 0.01 mm^−1^ at 532 nm. DIS system setup and measurement procedure are described elsewhere^24^.

#### Phantom mould and inclusions

For phantom studies, a versatile, modular phantom mould was created by 3D-printing (Figure 2). Phantom moulds were designed in Autodesk Fusion 360 (San Rafael, CA, USA) and printed using an Anet A6 Printer and polylactic acid (PLA PRO 1.75 mm PLA 3D Printer Filament 832-0232 (Yellow) and 832-0223 (White), RS Components, Corby, UK) as a base material. The phantom mould consists of two modules (Figure 2A). The inner module acts as the actual testing module and allows the measurement of specific testing parameters. The outer module functions as a frame for the inner module and prevents leaking of the bulk material and enables a firm positioning of the phantom. Indentations in the corners of the outer module allow insertion of structural support blocks to control the distance between detector and phantom surface. Customized handles can be inserted on diagonal sides of the inner module to facilitate the insertion and removal of the inner module from the outer module. For this work, two inner modules have been developed. The first inner module, the string module (Figure 2B), allows for the examination of basic technical parameters such as the sensitivity of the system to angularity or penetration depth through insertion of targets at different depths and angles. The second module, the tubing module (Figure 2C), permits the analysis of liquid contrast agents by including tubing (Fine Bore Polythene Tubing 0.58 mm inner diameter, 0.96 mm outer diameter, Portex) at different depths and directions.

**Figure 2:**
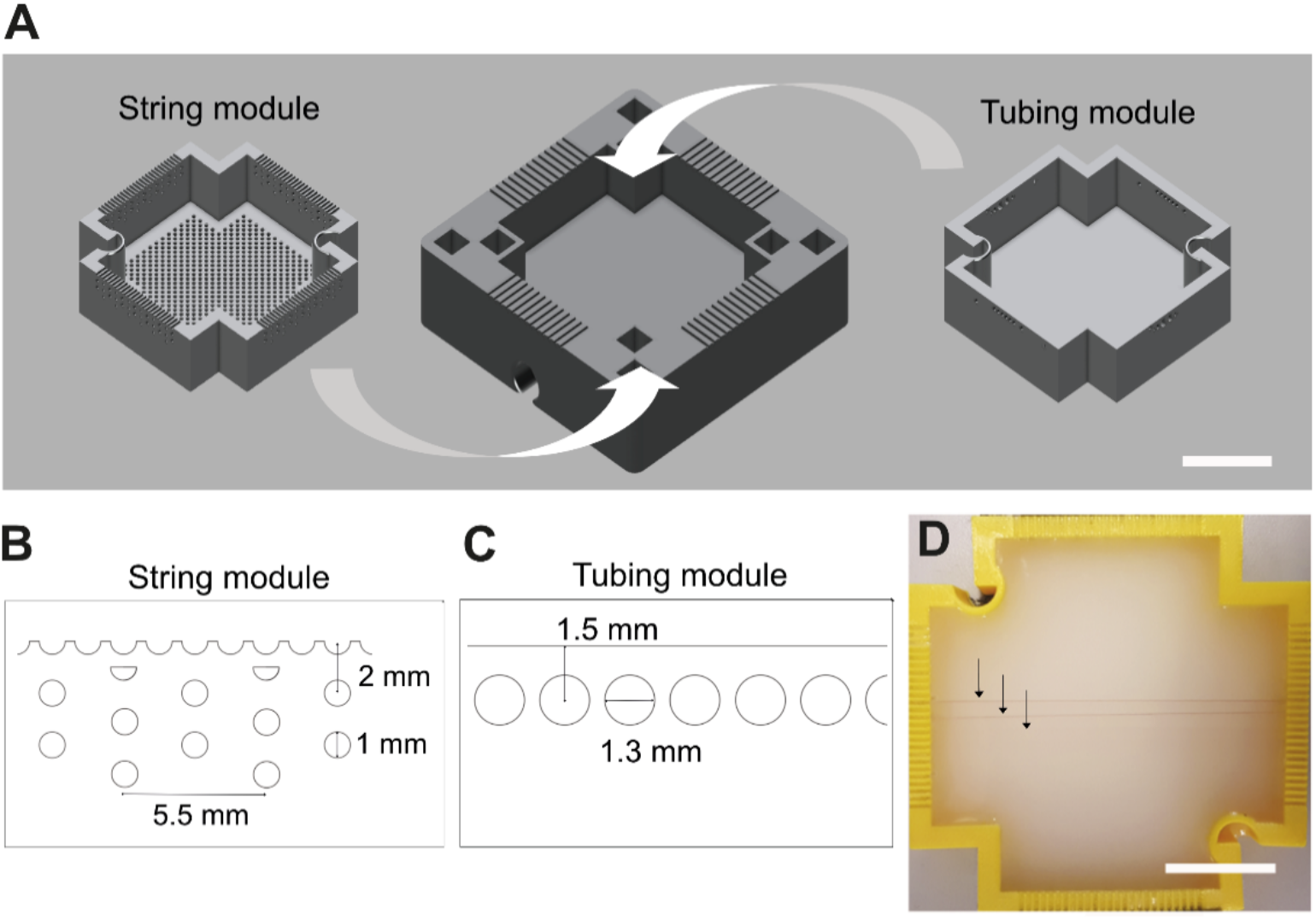
Phantom design for the technical validation studies. (A) The computer-aided designs of the outer (middle) and inner (right, left) modules used for this study are shown. Close-up side view of the image quality phantom modules: (B) String array allowing penetration depth and angular studies; (C) Tubing array allowing sensitivity studies. (D) Photograph of the 3D-printed string phantom module filled with agar and featuring targets at different depths (black arrows) is shown. Scalebars = 20 mm.

For evaluation of imaging artefacts, a dilution series of red ink (Cranfield Colours, Wales, UK) was used in the tubing module. For geometric sensitivity studies, the string module was deployed with red-coloured synthetic fibres (Smilco, Houston, TX, USA). These were chosen as imaging targets for the string module due to their similar size^26,27^ to murine vessels and high absorption at 532 nm.

### Animal handling

Procedures on small animals were performed under the authority of project (PE12C2B96) and personal (IA70F0365, I544913B4) licenses issued by the Home Office, UK. Studies were approved by the Cancer Research UK Cambridge Institute local animal welfare and ethical review bodies under compliance form numbers: CFSB2112, CFSB1567 and CFSB1745. All mice were housed in Tecniplast Green Line individually ventilated cages with APB6 bedding on a 12-h on/off light/dark cycle (7AM to 7PM) with 5R58 diet (PicoLab).

To evaluate precision of the system in imaging tumour-bearing mice, a cell line xenograft model and four patient-derived xenograft (PDX) models were used (Table 1). For the cell line model, subcutaneous tumours were established in male BALB/c nude mice (Charles River, age 8-10 weeks). 1.5×10^6^ PC3 prostate adenocarcinoma cells suspended in a mixture of 50 μL PBS and 50 μL matrigel (354248; Corning) were inoculated subcutaneously in both lower flanks of n=5 mice (resulting in n=10 tumours). For the PDX models, two luminal B patient-derived xenograft (PDX) models (AB580, STG143) and two basal PDXs (STG139, STG321) were implanted subcutaneously into the flank of 6-9 week-old NOD SCID gamma (NSG) mice (Jax Stock #005557) using the standard protocols of the Caldas laboratory biobank at the Cancer Research UK Cambridge Institute^28^ (n_AB580_ =12, n_STG143_ =10; n_STG139_ =26, n_STG321_=18). Before surgical implantation, cryopreserved breast patient-derived xenograft tumour fragments (~2 mm^3^) in freezing media (foetal bovine serum, heat-activated Thermo Fisher Scientific 10500064 +10% dimethyl sulfoxide Sigma D2650) were defrosted at 37°C, washed with Dulbecco’s modified eagle’s medium (Gibco 41966) and mixed with matrigel (Corning 354262).

**Table 1:** Overview of the mouse models used in the individual studies.

| Study | Model |
| --- | --- |
| Tissue type (healthy vs pathological) | STG321 |
| Operator (experienced vs unexperienced) | PC3, STG321, AB580 |
| Positioning (dorsal vs lateral) | AB580, STG143, STG139, STG321 |

For *in vivo* imaging, mice were anaesthetised using 3% isoflurane delivered in 50% oxygen and 50% medical air. If needed, the hair was removed around the area to be imaged by shaving and application of commercial hair removal cream. Mice were placed on a heat-pad maintained at 37°C inside the system chamber. Respiratory rate was maintained between 70-80 bpm using isoflurane (~1.5-2% concentration) throughout image acquisition.

### Image and statistical analysis

Imaging data was reconstructed using a beam-forming algorithm, which models the sensitivity field of the focused detector and generates 3-dimensional images^30–32^. The reconstructed images were analysed using MATLAB (v2020, MathWorks, Natick, MA, USA) and Fiji (v2.1.0)^33^. Statistical and regression analysis was performed using Prism (v9, Graphpad Software, San Diego, CA, USA). All data are shown as mean ± standard deviation (SD) unless otherwise stated. Coefficients of variation (COV) were calculated as the ratio of the SD to the mean, expressed as percentage. For multiple comparisons, one-way ANOVA followed by Tukey’s test was conducted. Correlation analysis between dorsal and lateral imaging positions was conducted using Spearman’s correlation coefficient due to non-normal data distribution.

#### Phantoms

For phantom image analysis, a fixed-sized rectangular region of interest spanning the length of the string was placed around each string within the image. For each string, the line profiles perpendicular to the string were extracted within the region of interest. The background was subtracted and a gaussian curve was fitted to the signal. The full width at half maximum (FWHM) and mean intensity value were extracted from the fit for each line profile. Subsequently, the mean and SD of all FWHMs and signal intensities of all line profiles were calculated to achieve final values for each string.

#### In vivo

For the *in vivo* repeatability studies, the blood volume was chosen as a comparison metric, as it is a commonly used variable of interest in preclinical studies, and sometimes used as a precursor for further downstream analysis of morphological features^34^. An existing pipeline^34^ was used to quantify the results. Briefly, image data was filtered in the Fourier domain in the XY plane to remove reflection lines, before being reconstructed using a backprojection algorithm in viewRSOM software (v2.3.5.2 iThera Medical GmbH, Germany) with motion correction and a voxel size of 20 × 20 × 4 μm^3^ (X,Y,Z). Reconstructed images were subjected to a high-pass filter^20^ to remove echo noise, followed by a Wiener filter to remove stochastic noise. Afterwards, a slice-wise rolling ball background correction^35^ was performed to achieve a homogenous background intensity. Segmentation was performed using a random forest classifier (ilastik v1.3.3^36^). For the classifier, n=20 *in vivo* images were used for training and n=14 *in vivo* images were used for testing. Then, all segmented images were passed through a 3D median filter to smooth and remove impulse noises.

## Results

### Phantom studies illustrate illumination, shadow and reflection artefacts in photoacoustic mesoscopy

To initiate our characterisation of photoacoustic mesoscopy performance, we first sought to investigate dominant artefacts present in the images. Artefacts emerging from absorbing vascular structures can degrade image contrast and confound interpretation. While clutter artefacts^37–39^ can be observed in the image background, affecting the overall imaging signal-to-noise ratio, three main classes of artefact can be seen to directly affect vessel analysis (Figure 3).

**Figure 3:**
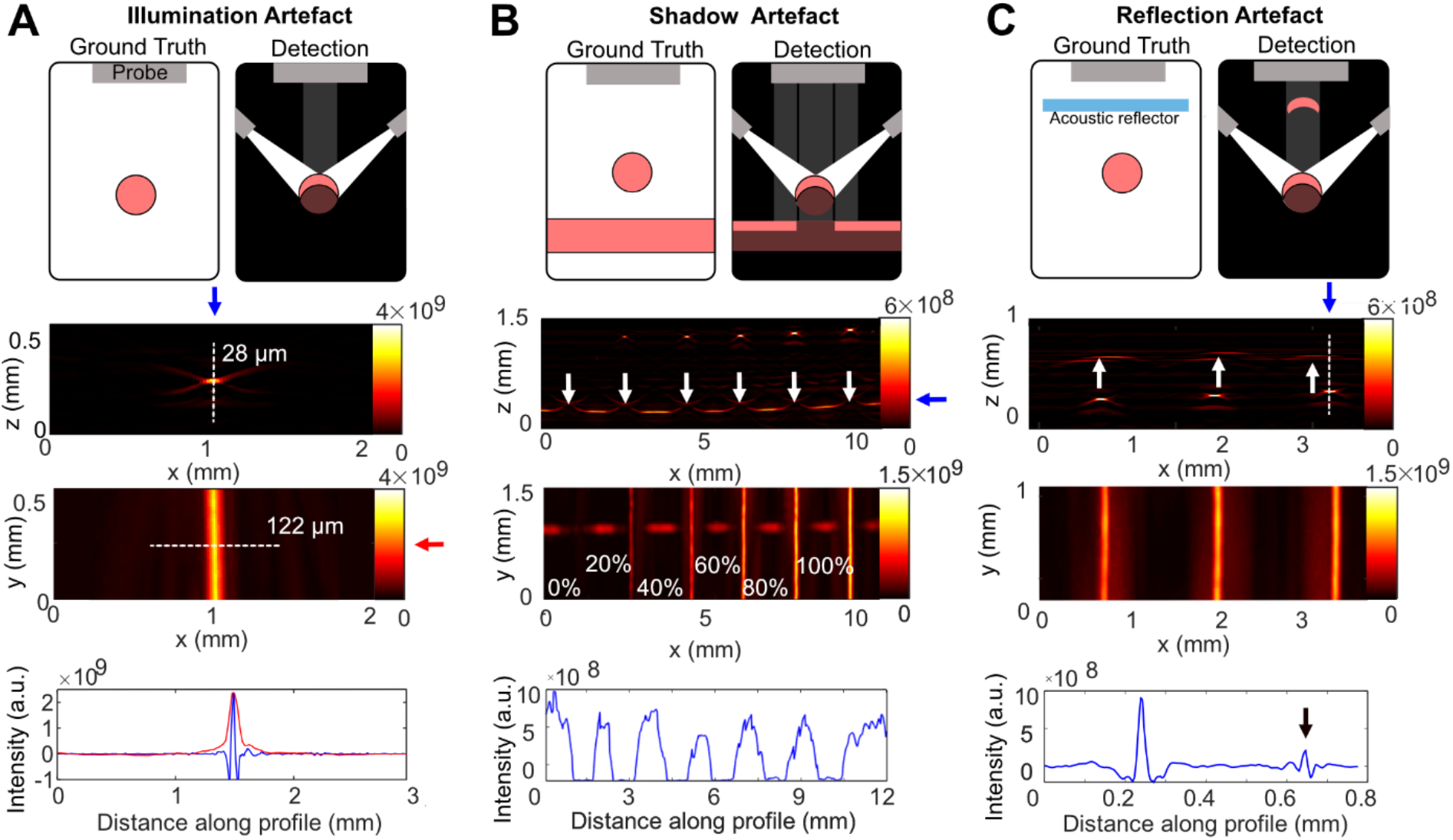
Overview of artefacts arising in the photoacoustic mesoscopy system. Explanatory schematics (first row), XZ MIPs (second row), XY MIPs (third row), and line profiles (fourth row) for the respective white dotted lines in the RSOM images for: (A) illumination artefact; (B) shadow artefact; and (C) reflection artefact. In A, a single string is shown, whilst in B, and C dilutions of red ink in tubing are displayed. In B, a tube of absorber is positioned perpendicular and beneath tubes of varying relative concentrations (up to 100%). The underlying tubing has the same concentration as the overlaying tubing with the highest concentration (100%). In C, the agar phantom/ultrasound gel interface acts as an acoustic reflector. White arrows depict the respective artefact; blue and red arrows indicate the respective line profiles plotted in the last row; the black arrow depicts the reflection artefact in the line profile.

First, illumination artefacts arise due to the limited top illumination of the field of view, leading to optical excitation and acoustic wave generation only in the upper part of absorbing structures that face the illumination and transducer array (Figure 3A). This results in inaccurate diameter estimations in the XZ plane compared to the XY plane^34^. For example, in the string phantom module, where the true diameter of each string is d_act_= 126 μm, the measurements from the reconstructed images agree only in the XY plane (d_xy_ =122 ± 7.8 μm) and substantially underestimate the value in the XZ plane (d_xz_=27 ± 3.2 μm). Second, shadow artefacts arise from obscuring objects, causing a signal loss in underlying objects due to strong optical or acoustic attenuation of the overlaying structure (Figure 3B). Shadow artefacts in this case are created by attenuation of the acoustic waves by the overlaying tubing, rather than by optical attenuation, as the length of the signal gap is independent of the signal intensity in the overlying tubes. Third, reflection artefacts occur due to presence of acoustic reflectors/scatterers or strong acoustic reverberations of the absorbing object itself^17,40,41^, leading to a signal echo near the object of interest (Figure 3C). They can occur in-plane or out-of-plane depending on the position of the absorber/reflector^42^. Awareness of these artefacts is of particular importance as illumination artefacts can lead to inaccuracies in quantification of vessel size, but more importantly, shadow and reflection artefacts can be mistaken for real structures in the image plane.

### The sensitivity of photoacoustic mesoscopy is orientation-dependent

We sought to systematically analyse how the geometric positioning of a vessel-like target in three-dimensional (3D) space affects the acquired signal. First, the impact of different target (string) depths on signal intensity and measured spatial dimensions was evaluated (Figure 4A,B). As expected, a significant signal loss occurred with increasing string depth (Figure 4C). Depth also impacted the quantified FWHM in XY direction with a significant decrease in measured FWHM with increasing depth (Figure 4D). Furthermore, the impact of vertical and horizontal target rotation on the acquired signal was tested. For testing of the horizontal rotation, a phantom with strings in a star-shaped pattern (Figure 4E,F) was imaged and the signal from each angled string quantified. To minimize inaccuracies in the target depth arising from the experimental preparation of stacking the strings, the phantom was rotated by 90° between image acquisitions. The angle of the string in relation to the direction of illumination was found to significantly impact the quantified mean PAI signal (Figure 4G), with the string in line with the two illumination fibres having the highest signal intensity. Encouragingly, there was no significant difference in the calculated FWHM between 90° and 0° or in the strings between the rotation steps (Figure 4H). For the vertical angles, a clear signal decay was found with increasing angle to the phantom surface (Figure 4 I-K). At 24°, the string could no longer be detected in the tissue-mimicking phantoms (Figure 4K). The size of the FWHM quantified from the MIPs remained stable for angles up to 20°, but beyond that it increased in value and variability due to the lower signal (Figure 4L). These results highlight depth- and angle-related limitations of the system resulting from the limited aperture of the illumination and transducer array, affecting signal detection and quantification.

**Figure 4:**
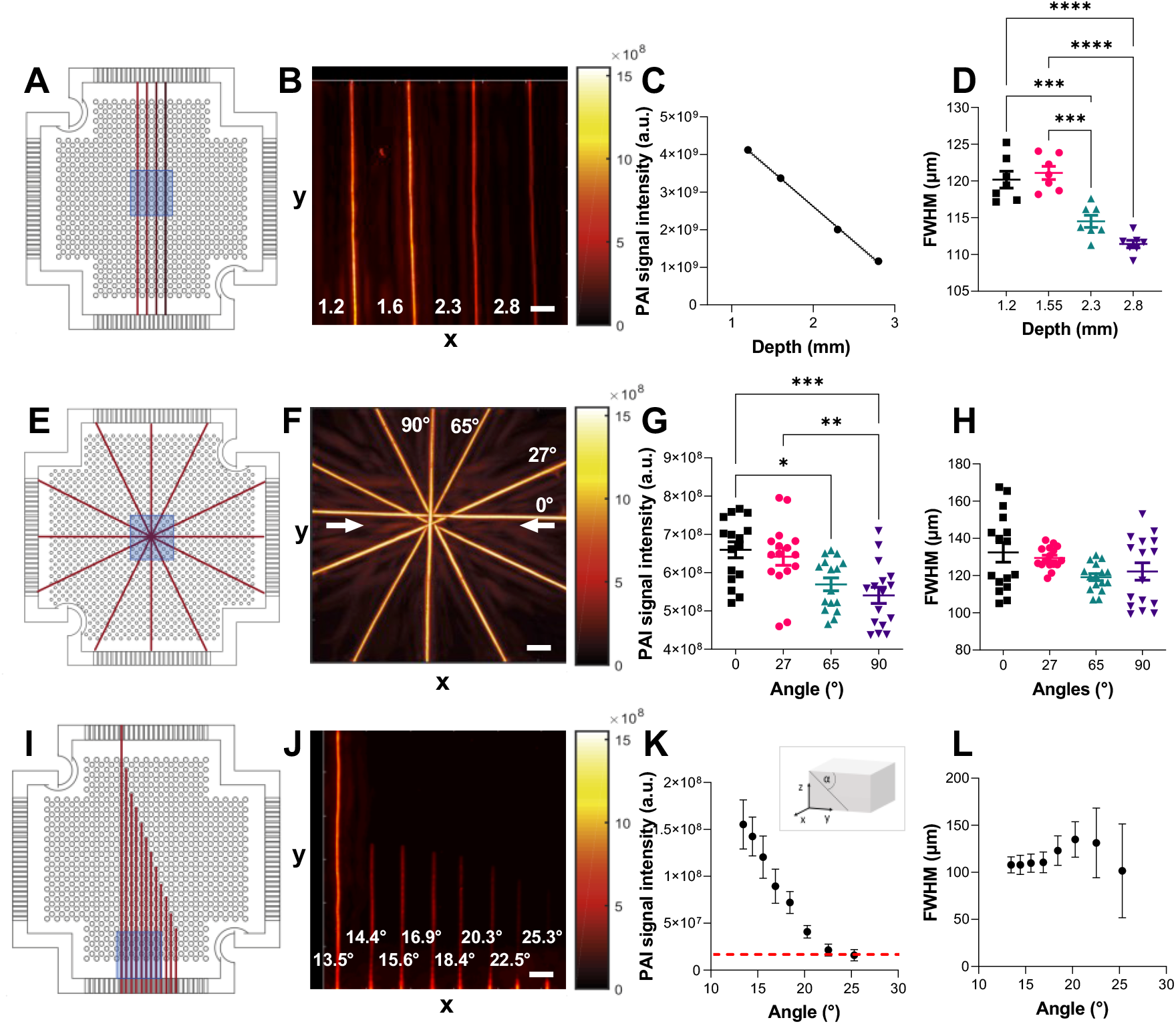
Geometric sensitivity of the photoacoustic mesoscopy system. Phantom configuration (first column), XY MIPs (second column), quantified signal intensity values (third column) and FWHM (fourth column) for: phantom with strings at different depths (A-D, n=3, R^2^=0.9991), phantom with horizontally angled strings (E-H, n=8), phantom with vertical angled strings (I-L, n=3). The field of view in the phantom configurations (corresponding to the MIPs) is marked in blue. The numbers in B depict the depth of the neighbouring string in mm. The direction of the optical fibres in F is marked with white arrows. Data displayed as mean ± SD. For figures D and G, significance was assigned using ANOVA (*p < 0.05, **p < 0.01, ***p < 0.001). Scale bars = 1.2 mm.

### Photoacoustic mesoscopy shows high precision for repeated measurement in phantoms

The signal stability was assessed through photoacoustic mesoscopy phantom measurements over time (Table 2, Figure 5). A mean COV of 1.2 ± 0.7 % (Figure 5A) was determined for repeated measurements in a single imaging session without repositioning of the phantom. With repositioning (removal from the system and immediate replacement), a slightly higher COV of 4.1 ± 2.4% was found (Figure 5B) for short-term studies (without switching off the system in between each runs), which increased to 9.6% for long-term studies (25 days, with switching off the system between each run). No slopes were significantly non-zero (without repositioning: p=0.1551; with repositioning [short term]: p=0.2858, with repositioning [long term]: p= 0.3955). The COVs were similar across different depths (Figure 5A,B), suggesting an excellent technical longitudinal repeatability for the system.

**Table 2:**
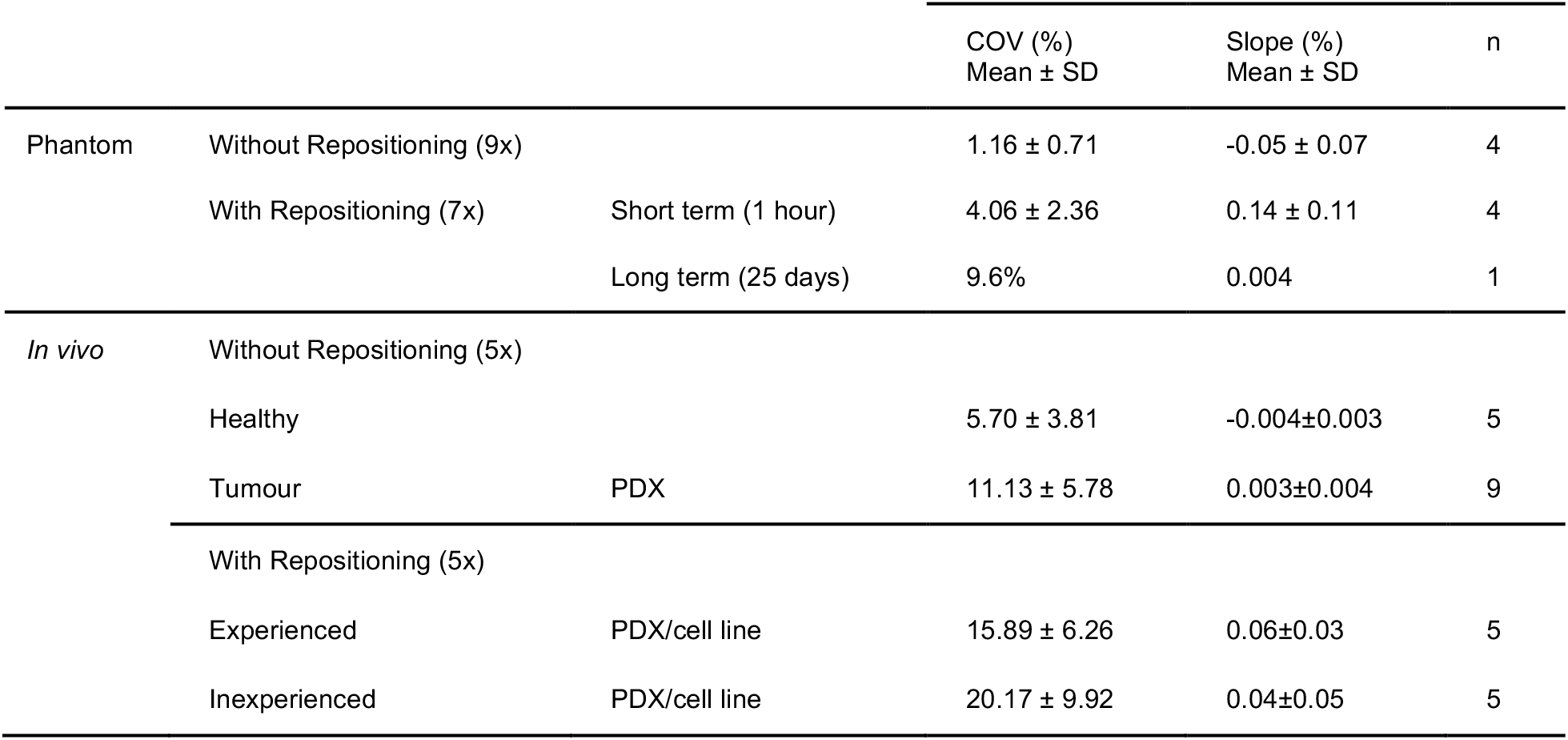
Coefficients of variation (COV) for temporal stability of the photoacoustic mesoscopy system. n refers to number of imaged mice/phantoms. PDX = Patient-derived xenograft.

**Figure 5:**
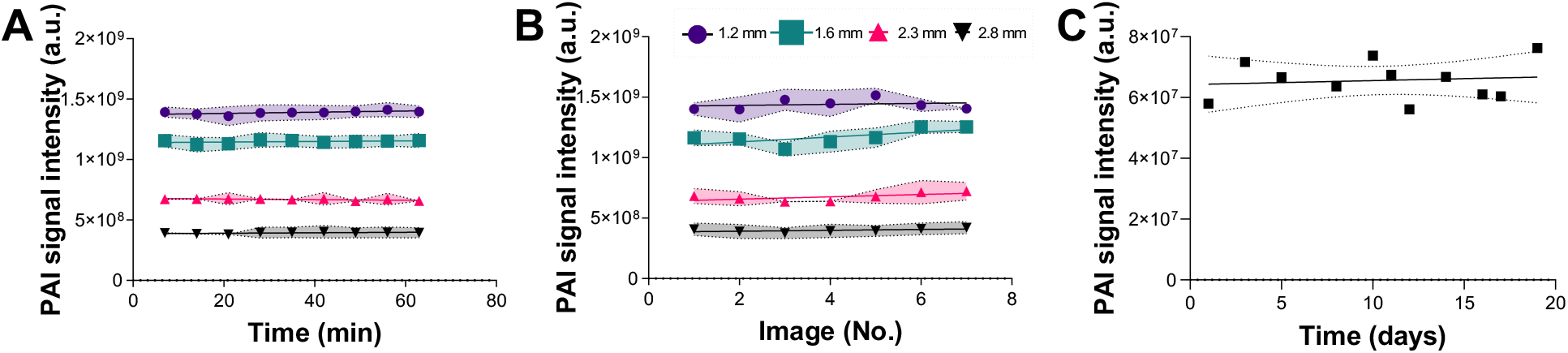
Temporal stability of the photoacoustic mesoscopy system. Signal stability in an agar phantom over time is shown along four strings embedded at four different depths (A) without replacement of the phantoms and (B) with replacement of the phantom between each sequential image acquisition. Legend indicates string depth. (C) Signal stability of a string embedded in a copolymer-in-oil phantom over a time frame of 20 days.

### Photoacoustic mesoscopy shows greater variation during *in vivo* application

The temporal stability of photoacoustic mesoscopy measurements was further evaluated *in vivo*, where additional sources of variation can arise (Figure 6A). Contributions from motion were minimised by appropriate positioning of the animals within the mouse bed and by application of a motion correction algorithm during image reconstruction. Optimal image acquisition in photoacoustic mesoscopy depends on a variety of factors, such as (1) adequate coupling of the of the tissue of interest to the transducer interface, (2) intactness of the coupling foil, and (3) purity of the coupling medium. Importantly, compression of the transducer interface on the target tissue must be avoided as it can cause signal drop out by impeding blood flow (Figure 6B,C).

**Figure 6:**
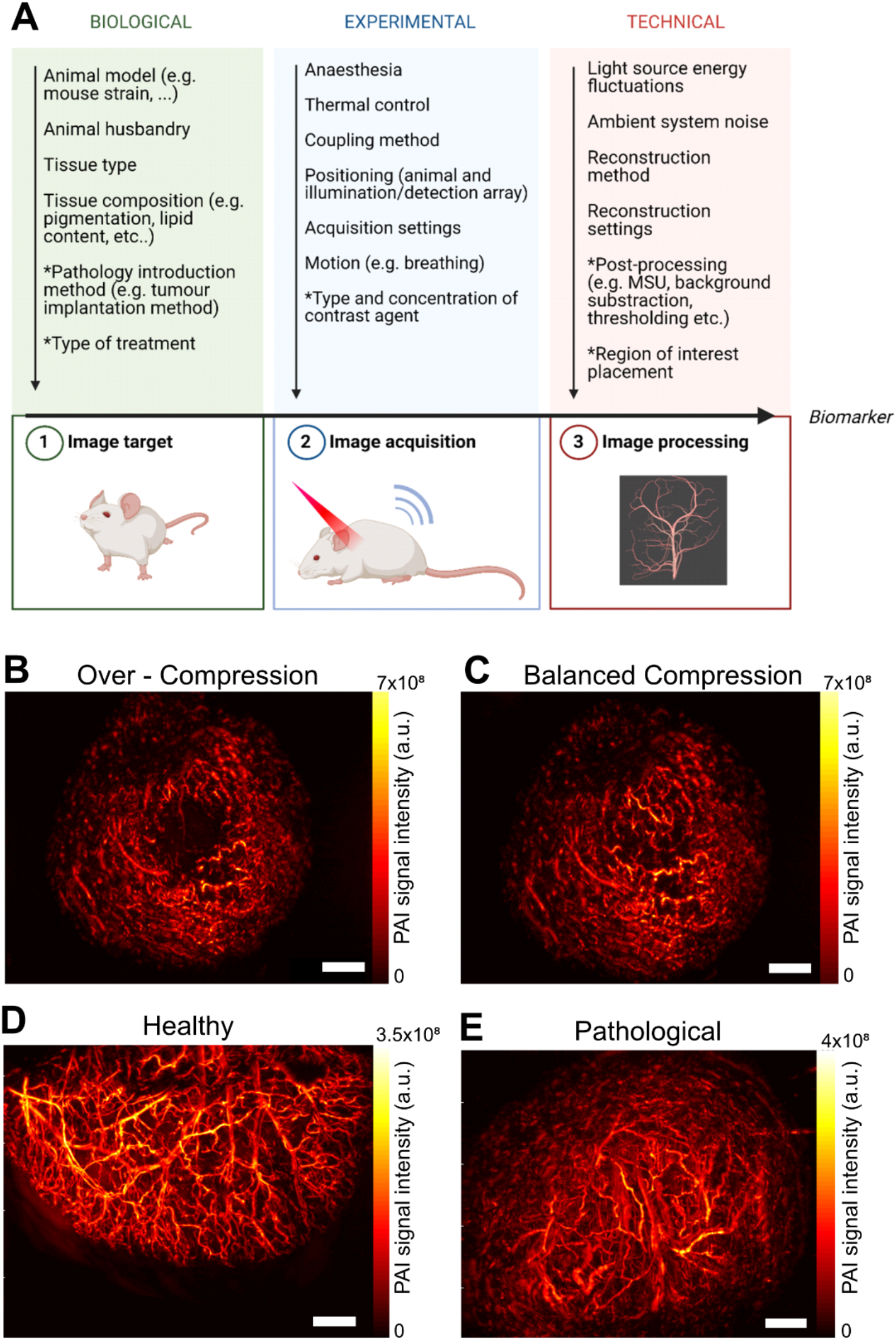
Biological and technical variation factors in photoacoustic imaging. A) Overview of biological and technical variation factors in photoacoustic imaging. MSU = Multispectral unmixing. Parameter marked with an asterisk describe variation factors that are only applicable in certain experiments/system types. Created with Biorender. B/C) Image acquisition in photoacoustic mesoscopy requires a balanced compression of the transducer interface on the tissue of interest. Whilst compression minimizes movement artefacts, it can also impact blood flow if applied too strongly, leading to signal loss (B). Balanced compression enables full visualization of all vessels (C). XY MIP of a PC3 tumour. Scale bar=1.5 mm. Representative mesoscopic XY MIPs for healthy (ear, D) and pathological (tumour, E) tissue are shown. Scale bars 1.5 mm.

For static imaging without repositioning *in vivo*, the blood volume in healthy tissue (Figure 6D) decreased over time (35 minutes; p=0.0065, Table 2, Figure 7A) with a mean COV of 5.7 ± 3.8 % (ear, Table 2). For pathological tissue (tumour, Table 2, Figure 7A), the mean COV increased to 11.1 ± 5.8 %, but no significant change over time (p=0.3622) could be observed.

**Figure 7:**
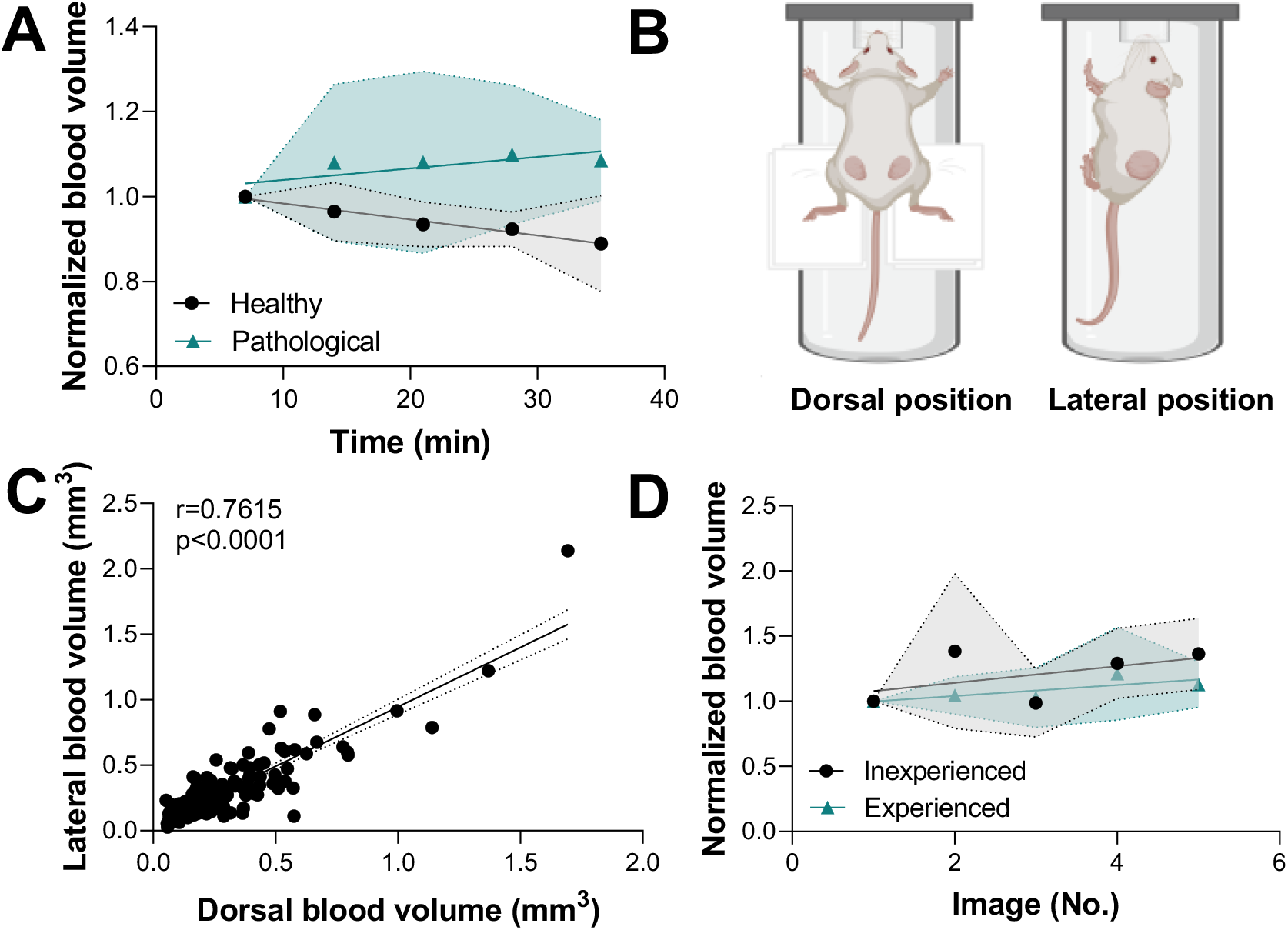
Impact of time, positioning, tissue and operator-dependent variation sources on the signal stability of the mesoscopic system in vivo. A) Blood volume is shown over time in healthy (black, n=6 ears, cell line model) and tumour tissue (green, n=9 tumours, PDX) without replacement of the mouse. Blood volume has been normalised by dividing each value with the first data point in the time series. B) Dorsal (left) and lateral (right) positioning of the mouse are indicated (figure created with Biorender). C) Correlation of calculated blood volume between dorsal and lateral positioning of the mouse is shown (n=161, PDX model, Spearman r=0.7615, R^2^=0.7687). D) Blood volume is shown in tumour tissue with replacement of the mouse by two different operators (each n=5, inexperienced operator=black, experienced operator=green, PDX and cell line models). Blood volume has been normalised by dividing each value with the first data point in the time series. All data is shown as mean ± SD.

The impact of the relative mouse positioning on the quantified blood volume was assessed for tumour imaging (Figure 7C). In our studies, the tumour position on the flank (a common location for preclinical oncology) allows for several different methods of animal positioning to be used, which will lead to different sub-volumes of the tumour being captured by the imaging system. A significant correlation (r= 0.76, p<0.0001) in the quantified blood volume between dorsal and lateral positioning of the mouse was found (Figure 7D). For the remainder of the studies, the leg position was used. Here, the influence of operator experience was also evaluated. With repositioning of the mouse (full removal of the mouse from the heat pad including cleaning off the ultrasound gel), mean COVs of 15.9 ± 6.3 % and 20.2 ± 9.9% for the blood volume were found for an experienced (preclinical RSOM imaging experience > 1 year) and inexperienced (no significant preclinical RSOM imaging experience) user, respectively (Figure 6D, Table 2). Taken together, our *in vivo* results suggest that photoacoustic mesoscopy provides a robust quantification of vascular data once users are experienced with the system.

## Discussion

Photoacoustic mesoscopy has shown great potential to become a widely used tool in biomedical research and clinical healthcare, but technical validation is required to ensure reliable data interpretation, reproducibility of data acquisition, and comparability between different studies. Using custom phantoms and *in vivo* analyses, we systematically analyse the impact of different variation sources on precision and accuracy of photoacoustic mesoscopy.

First, various artefact types that commonly appear in photoacoustic mesoscopy studies were exemplified and quantified. Accurate representation of vascular lumens is fundamentally limited in the photoacoustic mesoscopy geometry: since vessels are only illuminated from the top, images do not depict the full vessel volume (illumination artefact); signal loss also occurs for high angles of vessels relative to the detector, at increasing depth, or due to the presence of overlying structures (shadow artefact). The limited detection bandwidth of 10-90 MHz of the transducer translates to accurate representation of structures that are sized 12-120 μm^43^. Furthermore, reflection artefacts can occur that may be mistaken for real vessel structures. While some studies have attempted to identify and remove reflection artefacts, approaches are elaborate, using additional ultrasound measurements^37,40^, tissue deformations^16^, localized vibrations using acoustic radiation force impulses^19,38^, multi-wavelength illumination^17^, or training of convolutional neural networks^41^. Nonetheless, the full range of photoacoustic mesoscopy artefacts cannot yet be automatically detected or corrected and as such, understanding the resulting limitations on the acquired data is of utmost importance to prevent misinterpretation.

Second, limitations of the system were analysed that arise from three geometric factors significantly affect the measured signal intensity: (1) depth, and (2) horizontal and (3) vertical rotations of targets. In the commercial system tested, light is delivered from two optical fibres spaced 180° apart around the US transducer, leading to different light fluence at the sides orthogonal to this plane. Whilst these effects are minimal, and did not affect the quantification of target size, they significantly impact any associated measurements of signal amplitude. Illumination from four optical fibres placed at 90° around the US transducer, such as in other setups^44^, could help to mitigate this problem, but would increase system cost. The limited aperture of the US transducer^8^ also enabled signal quantification only up to an angle of 24° in our phantom, meaning structures angled more steeply could be missed. The relative impact of these factors also depends on the optical and acoustic properties of the surrounding medium, with higher signal loss in more attenuating media. As all these factors can significantly decrease accuracy of a measured structure, they should be considered when positioning a target within the system, or quantifying and evaluating acquired data.

After assessing accuracy-related factors, the precision of the system was evaluated. The system was found to be characterized by a high signal stability in both short-term and long-term phantom studies. This is in line with other studies showing less than 10% variation in PAI systems^20,21^. The values reported here are higher than those found previously for a commercial tomography system by the same vendor^20^ (COV_MSOT_=2.8% vs COV_RSOM_=9.6%), which is likely due to the fact that the distance between the target and detector is manually controlled by the user in the present system, affecting the measured signal intensity. Moreover, due to the higher resolution of the system, inhomogeneities in the coupling media may have a more pronounced effect on signal quantification. Ensuring constant transducer-target distance, as well as clear, bubble-free coupling agents, intact coupling foil and, if working with phantoms, homogeneous base media, will minimize the impact of these variation factors.

Finally, sources of variation *in vivo* were investigated considering time, tissue type, positioning and operator-dependent sources. As expected, a higher variability was found *in vivo* compared to in phantoms due to additional variations arising from motion and changes in vessel perfusion. When comparing healthy and tumour tissue, a higher variability was found in tumour tissue, which can be explained by the chaotic, tortuous and leaky nature of tumour vasculature^45^. Interestingly, a significant decrease in blood volume was seen in healthy vasculature over time. Similar observations have been made before in murine spleen and kidney at the macroscopic scale in a tomographic PAI system^20^, owing to perfusion changes under anaesthesia. Here, the ear was chosen as the healthy tissue of interest, as it is easily accessible by the system and often used as a site of interest in PAI studies^46–48^. However, as a peripheral organ the ear is not directly heated by the heat pad during image acquisition, which may have led to temperature-related vasoconstriction and thereby reduction of measured blood volume over time. Future work should explore the impact of temperature and isoflurane-related sources in more detail. A high correlation was found between the quantified blood volumes measured at two different data acquisition positions, demonstrating robustness in assessment of specimen-specific blood parameters. Variability between different operators was assessed, with only slightly higher values for an unexperienced user, confirming observations from other studies on photoacoustic precision^21^. Taken together, these observations demonstrate good precision of photoacoustic mesoscopy measurements in preclinical oncology models, across time and with different user experience.

There are several general considerations for photoacoustic mesoscopy that lead to high image quality, including keeping the coupling media and foil clean, intact and free of bubbles. Sample and coupling media temperature should be matched and care should be taken to maintain a constant transducer-target distance across repeated imaging sessions (e.g., by aligning the target of interest to the ‘focus-line’ of the transducer in the pre-scan). As target intensities are impacted by vertical and horizontal rotation, and penetration depth, the same positioning of the imaging object with respect to the illumination and detection plane should be used. For optimal quantification, the target of interest should be as close to the scan head and as horizontal relative to the scan head as possible and it should be noted that accurate dimensions can only be measured in x and y. For *in vivo* studies, in addition to the usual considerations of hair removal, maintaining breathing rate and animal body temperature, and minimizing motion, for mesoscopy measurements the transducer should be adjusted to avoid over- or under-compression.

Several limitations remain which should be addressed in future work. Inter-scanner variability has not been assessed as part of this study due to the logistical challenges of such a study. Similarly, the impact of motion correction, image reconstruction algorithms and data post-processing has not been investigated. First steps towards analysis of these computational variation sources in mesoscopic imaging have been taken elsewhere^34^, but future work should explore these technical factors of variability in more depth for further user guidance. With this work, we hope to guide data acquisition for similar PAI setups, thereby maximizing information retrieval and accuracy for future preclinical and clinical PAI studies.

## Acknowledgements

LH, ELB, PWS, TLL, and SEB acknowledge support from Cancer Research UK under grant numbers C14303/A17197, C9545/A29580, C47594/A16267, C197/A16465, C47594/A29448, and Cancer Research UK RadNet Cambridge under the grant number C17918/A28870. PWS acknowledges the support of the Wellcome Trust and University of Cambridge through an Interdisciplinary Fellowship under grant number 204845/Z/16/Z. TLL is supported by the Cambridge Trust. LH was funded from NPL’s MedAccel programme financed by the Department for Business, Energy and Industrial Strategy’s Industrial Strategy Challenge Fund. We thank the Cancer Research UK Cambridge Institute Imaging Core, Biological Resources Unit, Light Microscopy Core, and Research Instrumentation and Cell Services for their support in conducting this research. We also wish to thank Prof. Carlos Caldas, Dr Alejandra Bruna and Dr Wendy Greenwood for providing PDX tissue from their biobank at the CRUK Cambridge Institute and for assisting in the establishment of a sub-biobank that contributed the *in vivo* data presented in this manuscript.

## References

1. Omar, M., Aguirre, J. & Ntziachristos, V. Optoacoustic mesoscopy for biomedicine. Nat. Biomed. Eng. 3, 354–370 (2019).

2. Aguirre, J. et al. Precision assessment of label-free psoriasis biomarkers with ultra-broadband optoacoustic mesoscopy. Nat. Biomed. Eng. 1, 0068 (2017).

3. Taruttis, A. & Ntziachristos, V. Advances in real-time multispectral optoacoustic imaging and its applications. Nat. Photonics 9, 219–227 (2015).

4. Tanghetti MD, E. & Jennings, J. A comparative study with a 755 nm picosecond Alexandrite laser with a diffractive lens array and a 532 nm/1064 nm Nd:YAG with a holographic optic. Lasers Surg. Med. 50, 37–44 (2018).

5. Schwarz, M., Buehler, A., Aguirre, J. & Ntziachristos, V. Three-dimensional multispectral optoacoustic mesoscopy reveals melanin and blood oxygenation in human skin *in vivo.* J. Biophotonics 9, 55–60 (2016).

6. Haedicke, K. et al. High-resolution optoacoustic imaging of tissue responses to vascular-targeted therapies. Nat. Biomed. Eng. 4, 286–297 (2020).

7. Omar, M., Schwarz, M., Soliman, D., Symvoulidis, P. & Ntziachristos, V. Pushing the Optical Imaging Limits of Cancer with Multi-Frequency-Band Raster-Scan Optoacoustic Mesoscopy (RSOM). Neoplasia 17, 208–214 (2015).

8. Omar, M. et al. Optical imaging of post-embryonic zebrafish using multi orientation raster scan optoacoustic mesoscopy. Light Sci. Appl. 6, e16186–e16186 (2017).

9. Knieling, F. et al. Raster-Scanning Optoacoustic Mesoscopy for Gastrointestinal Imaging at High Resolution. Gastroenterology 154, 807–809.e3 (2018).

10. Aguirre, J. et al. Broadband mesoscopic optoacoustic tomography reveals skin layers. Opt. Lett. 39, 6297 (2014).

11. Schwarz, M., Omar, M., Buehler, A., Aguirre, J. & Ntziachristos, V. Implications of Ultrasound Frequency in Optoacoustic Mesoscopy of the Skin. IEEE Trans. Med. Imaging 34, 672–677 (2015).

12. Berezhnoi, A. et al. Assessing hyperthermia-induced vasodilation in human skin in vivo using optoacoustic mesoscopy. J. Biophotonics 11, e201700359 (2018).

13. Yew, Y. W. et al. Investigation of morphological, vascular and biochemical changes in the skin of an atopic dermatitis (AD) patient in response to dupilumab using raster scanning optoacoustic mesoscopy (RSOM) and handheld confocal Raman spectroscopy (CRS). Journal of Dermatological Science vol. 95 123–125 (2019).

14. Li, G. et al. Tripling the detection view of high-frequency linear-array-based photoacoustic computed tomography by using two planar acoustic reflectors. Quant. Imaging Med. Surg. 5, 57–62 (2015).

15. Li, M. et al. Linear array-based real-time photoacoustic imaging system with a compact coaxial excitation handheld probe for noninvasive sentinel lymph node mapping. Biomed. Opt. Express 9, 1408 (2018).

16. Jaeger, M., Siegenthaler, L., Kitz, M. & Frenz, M. Reduction of background in optoacoustic image sequences obtained under tissue deformation. J. Biomed. Opt. 14, 054011 (2009).

17. Nguyen, H. N. Y., Hussain, A. & Steenbergen, W. Reflection artifact identification in photoacoustic imaging using multi-wavelength excitation. Biomed. Opt. Express 9, 4613–4630 (2018).

18. Nguyen, H. N. Y. & Steenbergen, W. Three-dimensional view of out-of-plane artifacts in photoacoustic imaging using a laser-integrated linear-transducer-array probe. Photoacoustics 19, 100176 (2020).

19. Petrosyan, T., Theodorou, M., Bamber, J., Frenz, M. & Jaeger, M. Rapid scanning wide-field clutter elimination in epi-optoacoustic imaging using comb LOVIT. Photoacoustics 10, 20–30 (2018).

20. Joseph, J. et al. Evaluation of Precision in Optoacoustic Tomography for Preclinical Imaging in Living Subjects. J. Nucl. Med. 58, 807–814 (2017).

21. Wagner, A. L. et al. Precision of handheld multispectral optoacoustic tomography for muscle imaging. Photoacoustics 21, 100220 (2021).

22. Joseph, J., Ajith Singh, M. K., Sato, N. & Bohndiek, S. E. Technical validation studies of a dual-wavelength LED-based photoacoustic and ultrasound imaging system. Photoacoustics 22, 100267 (2021).

23. Chen, P. et al. Acoustic characterization of tissue-mimicking materials for ultrasound perfusion imaging research. Ultrasound Med. Biol. 48, 124–142 (2022).

24. Hacker, L. et al. A copolymer-in-oil tissue-mimicking material with tuneable acoustic and optical characteristics for photoacoustic imaging phantoms. IEEE Trans. Med. Imaging 1–1 (2021) doi:10.1109/TMI.2021.3090857.

25. Cabrelli, L. C. et al. Stable phantom materials for ultrasound and optical imaging. Phys. Med. Biol. 62, 432–447 (2017).

26. Müller, B. et al. High-resolution tomographic imaging of microvessels. Dev. X-Ray Tomogr. VI 7078, 70780B (2008).

27. Qian, B., Rudy, R. F., Cai, T. & Du, R. Cerebral artery diameter in inbred mice varies as a function of strain. Front. Neuroanat. 12, (2018).

28. Bruna, A. et al. A Biobank of Breast Cancer Explants with Preserved Intra-tumor Heterogeneity to Screen Anticancer Compounds. Cell 167, 260–274.e22 (2016).

29. Omar, M., Gateau, J. & Ntziachristos, V. Raster-scan optoacoustic mesoscopy in the 25–125 MHz range. Opt. Lett. 38, 2472 (2013).

30. Omar, M., Soliman, D., Gateau, J. & Ntziachristos, V. Ultrawideband reflection-mode optoacoustic mesoscopy. Opt. Lett. 39, 3911 (2014).

31. Schwarz, M. et al. Optoacoustic Dermoscopy of the Human Skin: Tuning Excitation Energy for Optimal Detection Bandwidth with Fast and Deep Imaging in vivo. IEEE Trans. Med. Imaging 36, 1287–1296 (2017).

32. Schwarz, M., Buehler, A. & Ntziachristos, V. Isotropic high resolution optoacoustic imaging with linear detector arrays in bi-directional scanning. J. Biophotonics 8, 60–70 (2015).

33. Schindelin, J. et al. Fiji: an open-source platform for biological-image analysis. Nat. Methods 9, 676–682 (2012).

34. Brown, E. L. et al. Quantification of Vascular Networks in Photoacoustic Mesoscopy. Photoacoustics 100357 (2022) doi:10.1016/j.pacs.2022.100357.

35. Sternberg, S. R. Biomedical Image Processing. Computer (Long. Beach. Calif). 16, 22–34 (1983).

36. Berg, S. et al. ilastik: interactive machine learning for (bio)image analysis. Nat. Methods 16, 1226–1232 (2019).

37. Schwab, H.-M., Beckmann, M. F. & Schmitz, G. Photoacoustic clutter reduction by inversion of a linear scatter model using plane wave ultrasound measurements. Biomed. Opt. Express 7, 1468 (2016).

38. Jaeger, M., Bamber, J. C. & Frenz, M. Clutter elimination for deep clinical optoacoustic imaging using localised vibration tagging (LOVIT). Photoacoustics 1, 19–29 (2013).

39. Wu, D., Tao, C. & Liu, X. Photoacoustic tomography extracted from speckle noise in acoustically inhomogeneous tissue. Opt. Express 21, 18061 (2013).

40. Kuniyil Ajith Singh, M. & Steenbergen, W. Photoacoustic-guided focused ultrasound (PAFUSion) for identifying reflection artifacts in photoacoustic imaging. Photoacoustics 3, 123–131 (2015).

41. Allman, D., Reiter, A. & Bell, M. A. L. Photoacoustic Source Detection and Reflection Artifact Removal Enabled by Deep Learning. IEEE Trans. Med. Imaging 37, 1464–1477 (2018).

42. Nguyen, H. N. Y. & Steenbergen, W. Reducing artifacts in photoacoustic imaging by using multi-wavelength excitation and transducer displacement. Biomed. Opt. Express 10, 3124 (2019).

43. IThera Medical. Instruction Manual RSOM Explorer Part III. 1.1, 8 (2018).

44. Wang, Y. et al. An Adjustable Dark-Field Acoustic-Resolution Photoacoustic Imaging System with Fiber Bundle-Based Illumination. Biosensors 11, 262 (2021).

45. Chung, A. S., Lee, J. & Ferrara, N. Targeting the tumour vasculature: insights from physiological angiogenesis. Nat. Rev. Cancer 10, 505–514 (2010).

46. Liu, C., Liang, Y. & Wang, L. Optical-resolution photoacoustic microscopy of oxygen saturation with nonlinear compensation. Biomed. Opt. Express 10, 3061 (2019).

47. Moothanchery, M. & Pramanik, M. Performance Characterization of a Switchable Acoustic Resolution and Optical Resolution Photoacoustic Microscopy System. Sensors 17, 357 (2017).

48. Lu, J. et al. In vivo photoacoustic imaging of blood vessels using a homodyne interferometer with zero-crossing triggering. J. Biomed. Opt. 22, 036002 (2017).

